# Ultrasensitive, Low-Input Detection of Avocado Sunblotch Viroid via RPA-CRISPR and Nanopore-Array Single-Bead Fluorescence Readout

**DOI:** 10.64898/2026.01.19.700023

**Authors:** Jiayi Xu, Xiaoqian Jiang, Mehdi Kamali Dashtarzhaneh, Yiding Zhong, Bhaskar Sharma, Ruonan Peng, Fatemeh Khodadadi, Ke Du, Chuanhua Duan

## Abstract

Rapid and sensitive detection of plant pathogens, such as the Avocado Sunblotch Viroid (ASBVd), is essential for early disease management and agricultural biosecurity. Yet, most current diagnostic methods not only require relatively large sample inputs but also often lack the ultrasensitivity required for reliable detection with scarce or minimally collected plant material. Here, we report a novel low-input but ultrasensitive diagnostic platform that integrates isothermal recombinase polymerase amplification (RPA), CRISPR-Cas12a detection, and a solid-state nanopore array for the detection of ASBVd. The system leverages CRISPR-Cas12a collateral cleavage activity to generate single-bead fluorescent signals, which are captured by a nanopore array through pressure-driven blockage. Our platform achieves a detection limit down to 1.68 copies/μL while using only 40 nL of bead-fluorophore mixture per readout, which is over 100-fold less than conventional assays based on fluorescent readout using an imaging reader, enabling detection from minimal avocado sample collection. We demonstrate robust binary classification of ASBVd-positive and -negative samples from multiple avocado tissue types and orchards in California. The assay requires just 60 minutes and operates entirely under isothermal conditions, avoiding the need for bulky PCR instruments and supporting on-site deployment with minimal equipment. This method provides a promising platform for field-deployable, ultrasensitive, and low-input diagnostics of viroids and other low-titer pathogens in plant or clinical settings.

## Introduction

Plant diseases pose a significant threat to global food security and agricultural sustainability, causing an estimated 20% to 40% reduction in crop yields worldwide^1^. Plant viruses are among the most economically damaging pathogens, causing a wide range of symptoms and substantial crop losses worldwide^2^. Plant viroids, in contrast, are the smallest known infectious agents, consisting of short circular RNA molecules without protein-coding capacity^3^. They pose particular challenges for crop production because they can cause severe symptoms while often remaining undetected in asymptomatic carriers. Accurate and early detection of both viruses and subviral pathogens such as viroids, is therefore critical for effective disease control, agricultural biosecurity, and propagation management.

These detection challenges are particularly acute in high-value crop systems such as avocado (*Persea americana Mill.)*, an increasingly important global commodity crop. Avocado sunblotch viroid (ASBVd), a viroid infecting avocado, causes severe damage including fruit deformation, tree dwarfing, and yield reduction^4^. Critically, symptomless carrier trees are considered as the main transmission source for ASBVd in orchards^5^, leading to substantial yield reductions ranging from 15% to 58% in Hass and Mendez trees^6^. This underscores the importance of early-stage detection for effective ASBVd management and prevention of its propagation through infected trees. However, reliable detection of ASBVd remains technically challenging due to the following primary factors. ASBVd is a small circular RNA molecule consisting of only 246-251 nucleotides^7^, which limits probe or primer design and amplifies inefficiencies, making detection difficult with standard molecular methods. Besides, the uneven distribution of the ASBVd within the host tree’s vascular system, particularly its low concentration and sporadic presence in leaves or fruits, presents a significant challenge for reliable sampling and early detection^8^. Furthermore, in asymptomatic or early-stage infections, viroid titers are often very low, necessitating the use of more precise and sensitive detection methods^5,9^.

The existing nucleic acid-based diagnostics remain limited in effectively detecting viroids. Over the past decades, early approaches such as polyacrylamide gel electrophoresis (PAGE) and dot blot hybridization have been employed to visualize or hybridize viroid RNA directly, offering qualitative results without amplification^10^. However, these methods suffer from low sensitivity, labor-intensive protocols, and limited applicability to low-titer infections^11,12^. Polymerase chain reaction (PCR)–based assays subsequently became the gold standard in viroid and virus diagnostics, with reverse transcription PCR (RT-PCR), quantitative PCR (qPCR), and droplet digital PCR (ddPCR) providing high sensitivity and enabling quantification of viral or viroid copy numbers at very low concentrations^13,14^. Despite their analytical performance, PCR-based methods require sophisticated laboratory infrastructure, trained personnel, and multi-step workflows involving complex thermal cycling. As such, they are generally restricted to centralized laboratories, limiting their applicability in field settings^15^. To address these limitations, isothermal amplification techniques such as loop-mediated isothermal amplification (LAMP) and recombinase polymerase amplification (RPA) have been developed to simplify reaction conditions and reduce instrument dependence^16–19^. LAMP offers robust amplification but demands complex primer sets and is prone to non-specific amplification^20,21^, whereas RPA operates at near-ambient temperatures (37–42 °C), requires simpler primers, and achieves rapid amplification within minutes, making it particularly attractive for portable diagnostics. In parallel, CRISPR-Cas–based diagnostics have emerged with remarkable specificity and single-molecule sensitivity, enabling nucleic acid detection via trans-cleavage of reporter probes^22,23^. Nevertheless, most implementations still rely on fluorescence or lateral flow readout equipment, which requires relatively large sample volumes, depends on bulk fluorescence measurements that can suffer from background interference, and often lacks sufficient sensitivity for low-titer pathogens^24^.

In this study, we present a novel detection system for ASBVd that integrates RPA, CRISPR-Cas12a–mediated signal activation, and nanopore array–based single-particle fluorescence analysis. After RNA extraction, reverse transcription, and DNA amplification, fluorescently labeled magnetic beads (MB-Cy5) are generated via Cas12a-triggered cleavage of quenched fluorescent probes. These beads are then quantified using a solid-state nanopore array, where each captured bead produces a localized fluorescent signal at a defined array site. Each readout requires only 40 nL of MB-Cy5 solution—several orders of magnitude less than the volumes typical of PCR or other conventional fluorometric approaches. This efficiency not only reduces reagent consumption but also allows detection from minimal starting material, facilitating early diagnosis even from limited samples. This approach enables ultrasensitive, semi-digital detection of ASBVd with a limit of detection down to 1.68 copies/μL (1 × 10L³ ng/μL total sample cDNA). Operating under isothermal conditions, our system eliminates the need for bulky thermal-cycle equipment and supports future field deployment using only simple heating sources and compact imaging modules. The platform holds promise for early viroid diagnosis in agriculture and could be extended to other low-abundance pathogens in future applications.

## Results and Discussion

### Assay Design and Sample Preparation

Figure 1a illustrates the sample processing pipeline from avocado tissue to cDNA template preparation. Avocado tissues were collected and subjected to total RNA extraction. Following purification, reverse transcription was performed to generate complementary DNA (cDNA) for use as the template in subsequent amplification reactions.

**Fig. 1:**
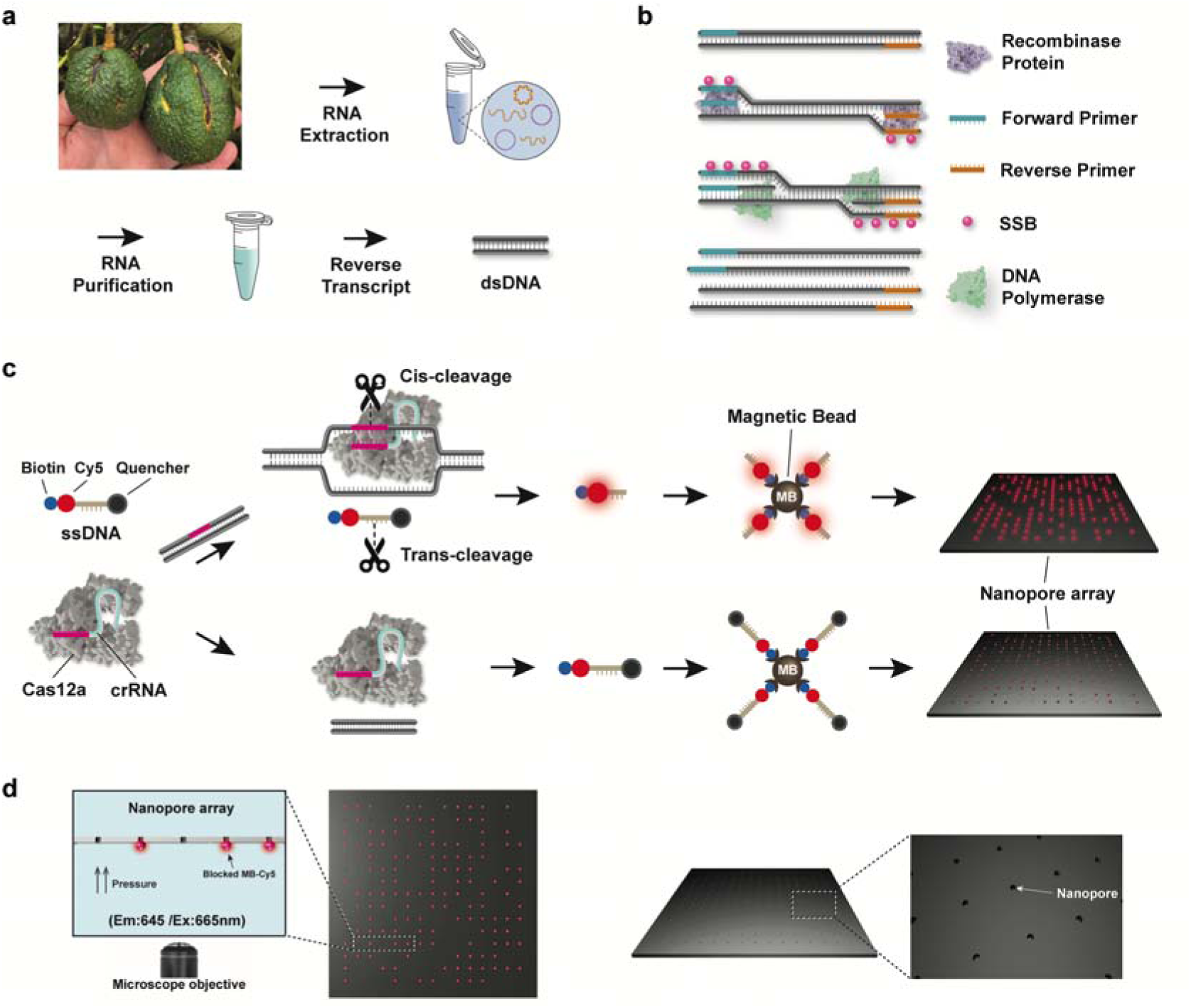
Schematic of RPA-CRISPR-based nanopore array fluorescence readout system. **a** Acquisition of total cDNA from infected Avocado tissue. **b** Mechanism of RPA amplification. Recombinases assist in primer binding to target DNA, while SSB proteins stabilize the strands. Polymerases then use building blocks (dNTPs) to form new DNA strands. **c** Mechanism of CRISPR-Cas12a detection and MB-Cy5 signaling on nanopore array. The fluorophore-quencher probe cleavages and Cy5 fluorophore is captured by magnetic beads (MBs) through biotin. MBs are then applied to the nanopore array. **d** Illustration of the fluorescent readout using blocked MBs.

The RPA mechanism is depicted in Fig. 1b, showing the coordinated action of recombinase proteins, single-strand DNA-binding proteins (SSB), and DNA polymerase. Recombinase proteins facilitated primer binding to target DNA sequences, while SSB proteins stabilized displaced strands. DNA polymerase then extended primers along the template, producing amplified products under isothermal conditions.

Figure 1c demonstrates the CRISPR-Cas12a detection mechanism. When the crRNA-Cas12a complex recognizes target DNA, cis-cleavage of the target sequence occurs, simultaneously triggering trans-cleavage of fluorescent ssDNA probes. This dual cleavage activity results in Cy5 fluorescent signal generation proportional to target presence. The cleaved products are subsequently conjugated with magnetic beads (MBs) to form MB-Cy5 complexes.

After the conjugation, the bead solution is diluted to 580 aM (0.5 mL total volume, corresponding to an original volume of only 40 nL) and loaded onto the *cis* side of the nanopore array, which is placed in a custom-made holder (see detailed description in Yazback *et al.* ’s publication^25^). Nanopores with a diameter of 400 nm were fabricated in a 15×15 array for this detection platform. A pressure difference of 0.5 atm was applied via a gas pressure regulator across the array to drive both unlabeled MB and MB-Cy5 toward the pores, capturing them individually at designated sites. After 1 min of pressure-assisted capture, the nanopore array was imaged by an inverted fluorescence microscope (*cis* side facing down). The design of the nanopore array and its supporting membrane enables quantification of the fluorescence of each captured MB while effectively minimizing background signals from other uncaptured MBs in the out-of-focus solution. The final readout of this platform is the fluorescent MBs ratio, representing the proportion of fluorescent-labeled MB-Cy5 among all captured MBs. This precise measurement approach provides high sensitivity for low-concentration fluorescent beads, enabling low-titer diagnostics from minimal sample input.

### Validation of Nanopore Array-Based Particle Detection

To evaluate the ability of the nanopore array platform to distinguish fluorescently labeled nano/micro beads, we first performed control experiments with non-biological samples. MB-Cy5 was used as the positive group, while unlabeled MBs served as the negative control. To simulate different detection scenarios, three bead mixtures containing 0%, 50%, and 100% MB-Cy5 were prepared, representing high, medium, and zero fluorescence conditions, respectively. Each mixture was diluted to a final concentration of 580 aM, and subjected to nanopore array blockage assays. Fluorescent images were taken after applying pressure to capture the beads at the nanopore array sites.

Scanning electron microscope (SEM) images of nanopore arrays before and after bead blockage are shown in Fig. 2a and 2b, respectively. It is clear from the SEM images that, after the blockage experiment, each captured MB was localized to a corresponding nanopore site and no multiple blockages at the same pore were observed - an important feature to avoid increased fluorescence from bead aggregation.

**Fig. 2:**
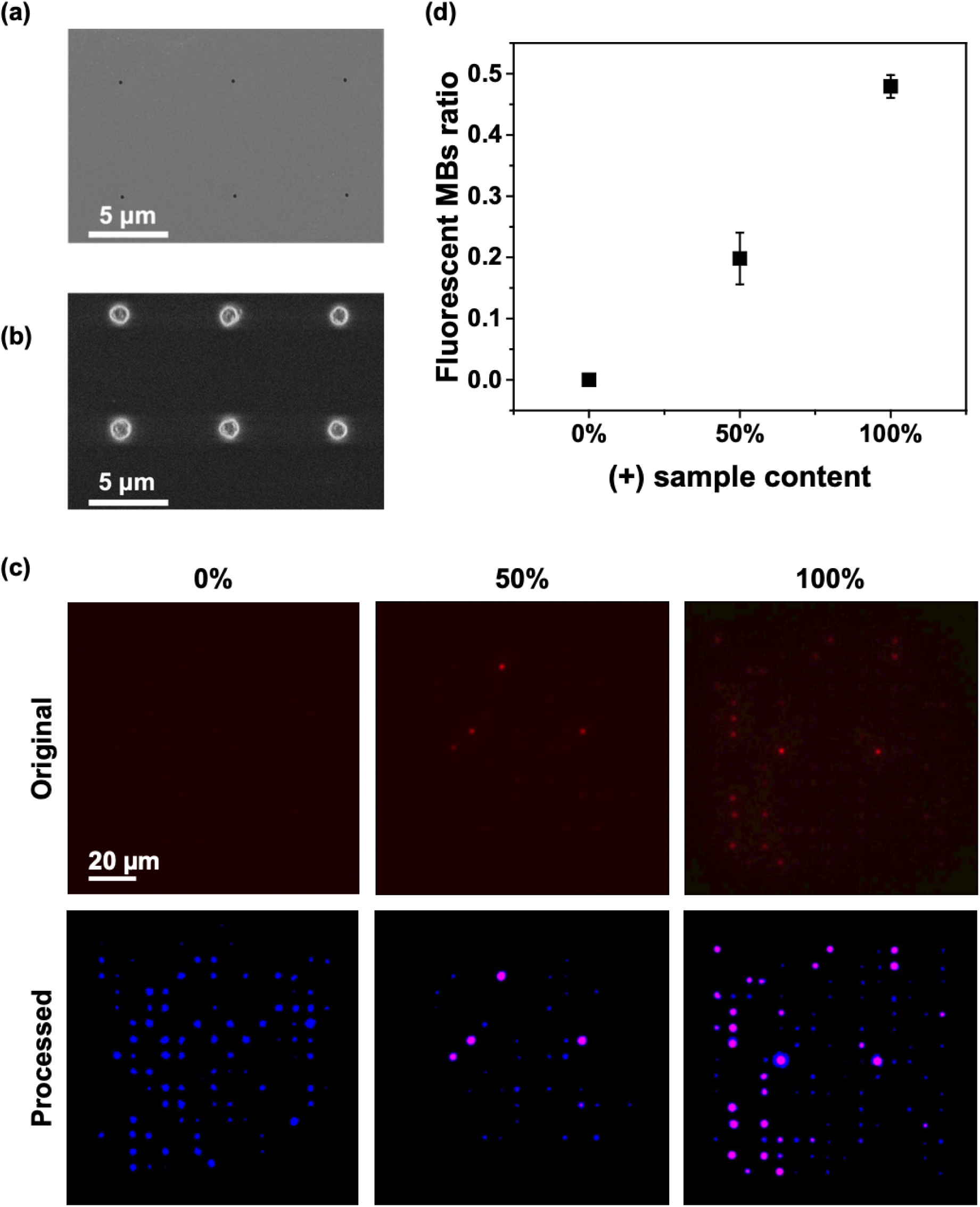
Nanopore array for MBs and MB-Cy5 fluorescence ratio detection. **a** Scanning electron microscope images before and **b** after MBs blocking experiments. **c** Fluorescent images and processed images for MB-Cy5 and MB mixtures, where the fluorescent beads are marked in pink in the processed images. **d** Fluorescent MBs ratio for different mixtures. Each condition is repeated on 3 different nanopore arrays.

Figure 2c shows the representative fluorescent image of the captured beads on the nanopore array. Only very faint fluorescence signals were detected in the 0% group (composed entirely of unmodified MBs), most likely due to background noise or the intrinsic autofluorescence of the magnetic beads. To eliminate such effects, a fluorescence intensity threshold was established across all three conditions to exclude weak background signals and reliably identify truly fluorescent MBs. As shown in Fig. 2c, regions above the threshold are highlighted in pink for fluorescent MBs, while beads below the threshold are displayed in blue. In the 100% positive group (MBs-Cy5), fluorescence intensity was heterogeneous, with some beads exhibiting markedly stronger signals than others. Fluorescent beads were also present in the 50% group, but the proportion of highly fluorescent MBs was clearly higher in the 100% group.

The ratios of fluorescent MBs calculated from three independent nanopore arrays are summarized in Fig. 2d. These ratios showed clear discrimination between groups, validating the ability of the nanopore array platform to distinguish varying levels of fluorescent labeling. It is worth noting that the fluorescent ratio observed in the 50% group was not exactly half of that in the 100% group. This discrepancy likely arises because only the MB-Cy5 population undergoes additional labeling and washing steps, during which a small portion of beads may be lost. Nevertheless, the observed trend confirms that the platform can effectively reflect relative proportions of fluorescently labeled beads. In real-sample detection, minor nonspecific loss of MB-Cy5 during washing and processing steps does not affect the fluorescent ratio, and thus does not compromise the performance or accuracy of the nanopore array readout.

This nanopore array platform enables directly imaging MBs in solution at attomolar concentrations. By physically capturing beads at defined, regularly spaced sites, the platform ensures single-particle isolation and precise fluorescence quantification without signal overlap or ambiguity. In addition to the fluorescent MBs ratio, we also evaluated the initial capture frequency—the rate of MB-Cy5 blockage events—as an alternative readout (Fig. S1). As previously described by Yazback *et al.*^25^, this dynamic information also correlates with fluorescent nanoparticle concentration. Our results showed a consistent trend as capture frequency decreased with lower proportions of MB-Cy5. However, capture frequency results showed higher variability across replicates. This is likely due to its strong dependence on controlled conditions, such as nanopore size uniformity, total bead concentration, sample purity, and potential residues from previous runs. In contrast, the fluorescence ratio proved more robust and reproducible under our experimental setup, since it only relies on the fluorescent portion of MBs. Furthermore, the fluorescence ratio requires only a single fluorescence imaging step after the capture process, which is much simpler than the initial frequency evaluation (which requires continuous fluorescence imaging of the nanopore array at the beginning of the capture process, posing challenges for the fluorescence camera). As a result, the nanopore array platform using the fluorescence ratio holds great promise for multiplexed sensing of different samples. It is worth noting that impurities from residual biomolecules and environmental contaminants may block a fraction of the nanopores, reducing the total number of analyzable MBs. This could introduce some variability in the measurements, especially compared to bulk methods like plate readers that analyze large populations of beads. However, each measurement consistently produced a reliable positive or negative result. Therefore, fluorescence ratio was adopted as the primary readout for this study. Having validated the quantitative readout capability of the nanopore array platform, we next applied it to the detection of ASBVd in field-collected avocado samples.

### ASBVd Identification in Real-World Avocado Samples

To evaluate the applicability of our detection system under real-world conditions, a diverse set of avocado tissue samples, such as leaves, flowers, and fruits, was collected from multiple orchards (Table S1). Following collection, total RNA was extracted from each sample and reverse-transcribed into cDNA. ddPCR assays were then conducted to identify the presence or absence of ASBVd in each specimen (Fig. S2). Based on the ddPCR results, seven ASBVd-positive samples from four different trees and eight ASBVd-negative samples from another set of four trees were selected to serve as representative inputs. The positive samples cover a wide range of ASBVd concentrations, from 2.4 to 1919 copies/μL.

For assay optimization, two CRISPR RNAs targeting ASBVd and their corresponding primer pairs (sequences listed in Table S2) were tested. The results are presented in Fig. 3a, with excitation and emission wavelengths set at 645 nm and 665 nm. The data demonstrate that proper DNA amplification by RPA significantly enhanced fluorescent signals in the CRISPR-Cas12a assay compared to reactions without RPA. While crRNA2 with primer set 2 generated higher overall fluorescence, crRNA1 with primer set 1 provided better discrimination between positive and negative samples. Based on this superior specificity, crRNA1 and primer set 1 were used in all subsequent evaluations.

**Fig. 3:**
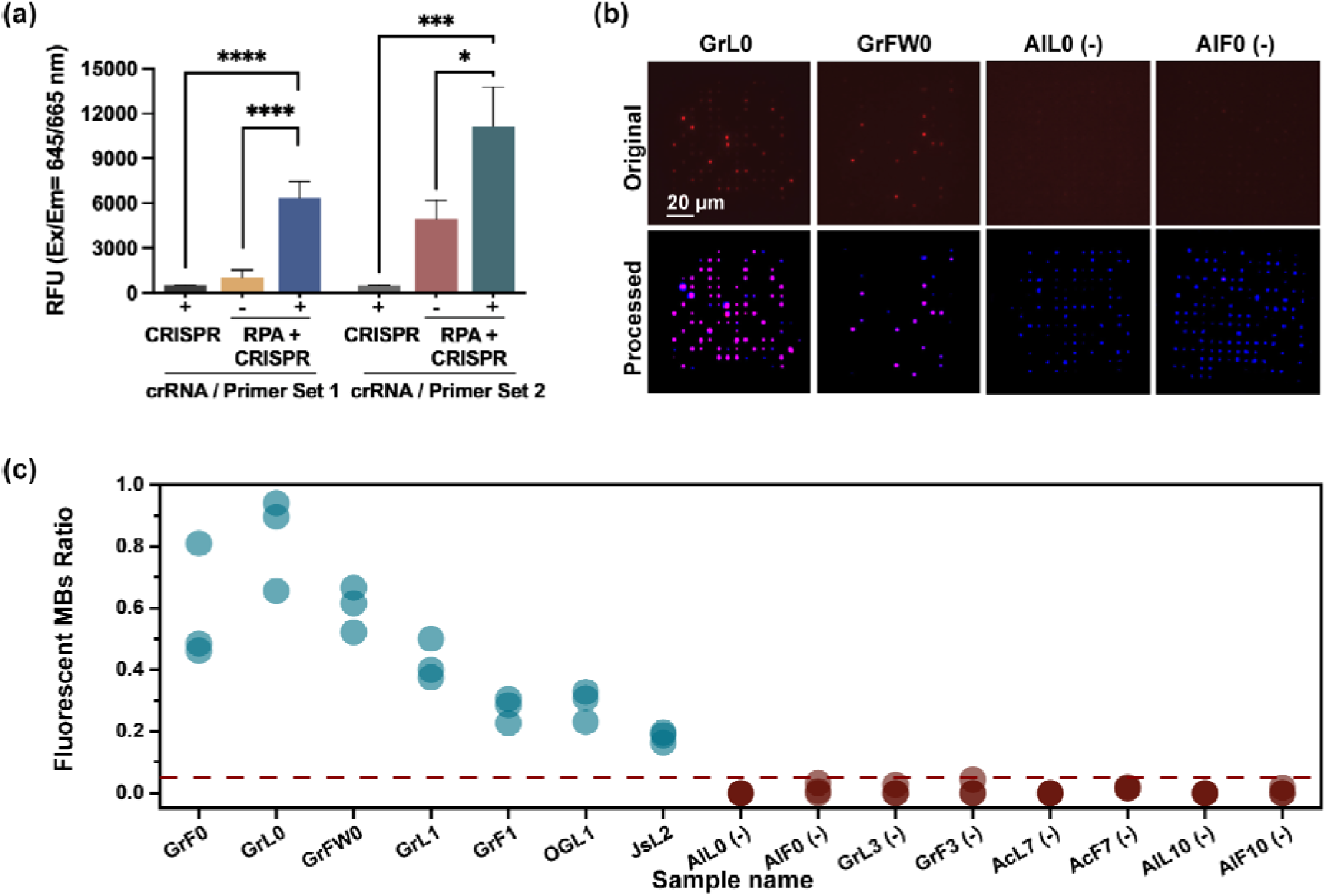
Evaluation of assay performance using different primer designs and different avocado tissues. **a** Selection of RPA primer and crRNA designs. **b** Representative images of nanopore array results for two positive samples on the left and negative samples on the right. For each sample, the original image in the first row shows the raw fluorescent image captured by fluorescence microscopy. The processed image displays only the identified MBs in blue, and fluorescent MB-Cy5 in pink against a black background. Each sample image not shown here is included in the Supplementary Information. **c** Detection results for all 15 different avocado samples. Each sample name represents the orchard (first letter), sample type (F, fruit; L, leaves; FW, flowers), and tree number. Detailed sample information is provided in Table S1. The first 7 samples are identified as positive by ddPCR, while the other 8 are negative (marked as (-) in the figure). Each condition is repeated on 3 different nanopore arrays.

Before binding with the CRISPR mixture, TEM analysis was performed to characterize the magnetic beads used in the detection system (Fig. S6). The beads exhibited appropriate morphology for biomolecular binding, confirming their suitability as solid-phase substrates for magnetic separation and fluorescence-based readout.

MBs derived from each of the 15 avocado tissue samples were analyzed by nanopore array blockage. Following one minute of pressure-driving loading, fluorescent images were captured under identical exposure conditions. Representative fluorescent images from two positive samples and two negative samples are shown in Fig. 3b. Positive groups exhibited clear fluorescent signals from MB-Cy5, whereas only faint signals were detected in the negative groups, suggesting the absence of target DNA.

Fluorescent ratios from all 15 samples are summarized in Fig. 3c. To quantitatively interpret the results, a fluorescence ratio threshold of 0.05 was established based on signal levels observed in negative controls. Applying this threshold, any sample exhibiting a fluorescent MBs ratio above 0.05 was classified as ASBVd-positive (in blue). Under this criterion, all ddPCR-confirmed positive samples were successfully identified by our nanopore-based platform, while all ddPCR-confirmed negatives remained below the threshold (in red), confirming the detection system’s robustness and consistency with established PCR diagnostics.

Beyond this binary classification, fluorescent ratios varied among positive samples. Higher-titer samples (e.g., GrF0, 1919 copies/μL and GrFW0, 704.3 copies/μL) generally yielded relatively higher ratios of fluorescent MBs, whereas the lowest-titer sample (JsL2, 2.4 copies/μL) showed the weakest. However, not all measured fluorescent ratios scale with ASBVd concentration. A similar non-linear trend was also observed in bulk fluorescence assays (Fig. S3). This likely reflects the multi-step nature of the RPA–CRISPR cascade, which does not yield a strictly proportional signal to template input. Although this non-linearity prevents direct quantification of template concentration, successful detection of the lowest-titer sample (JsL2, 2.4 copies/μL) highlights the practical sensitivity of our platform. Dashtarzhaneh *et al.* recently developed a digital LAMP (dLAMP) method for sensitive detection of ASBVd in California orchards^18^. However, some low-titer samples (JSL2 at 2.4 copies/μL) could not be consistently detected using the dLAMP method. Notably, our newly developed platform successfully generated a positive signal from the same JSL2 sample, highlighting its superior performance in ultra-low titer conditions.

In addition to its discrimination capability, the nanopore array detection platform also offers a potential advantage of operating with minimal sample volumes. Conventional fluorescence characterization by plate readers typically requires a 10-50 μL CRISPR reaction mixture (dependent on plate format) to ensure an adequate optical path for reliable detection. In contrast, the entire CRISPR-Cas12a reaction in our assay yields 20 µL of MB-Cy5 product, yet each nanopore array readout consumes only 40 nL. This drastic reduction means that large reaction volumes are not necessary for reliable detection, and the upstream amplification and CRISPR volume can therefore be reasonably reduced without compromising assay performance. As a result, the total amount of plant tissue required and the consumption of costly reagents can both be minimized, which is highly beneficial for field use where sample availability and preparation capacity may be constrained.

This low-input capability is enabled by the minimal sample volume required for the nanopore array detection platform, together with the RPA-CRISPR-based ASBVd to MB-Cy5 conversion. On one hand, the nanopore array provides single particle recognition requiring only a minimal number of target particles (on the order of 1 to 100) to achieve an accurate readout. On the other hand, our detection strategy incorporates a dual-modified single-stranded DNA probe engineered with both Cy5 fluorophore and biotin modifications. This bifunctional design served multiple critical roles: the Cy5 modification enables fluorescent signal generation for target detection, while the biotin moiety facilitates conjugation to streptavidin-coated magnetic beads. The magnetic bead coupling not only enhances target capture and purification efficiency through magnetic separation during washing steps, but also proves essential for the subsequent nanopore array-based visualization platform.

Together, the nanopore array detection platform not only achieves reliable ASBVd identification across diverse field samples, but also operates effectively with minimal sample input—an essential feature for resource-limited and on-site diagnostic applications.

### Sensitivity of the Detection Platform

To further validate the sensitivity and dynamic range of our detection system, we next performed amplification and CRISPR-mediated fluorescent labeling using a serial dilution of the GrF0 sample, where the total cDNA concentrations ranged from 1 to 10^-4^ ng/µL (1985 to 0.24 copies/µL, with ddPCR results shown in Fig. S4). The resulting fluorescently labeled MBs were analyzed using the same nanopore array blockage assay, and fluorescent ratios were calculated for each concentration group.

As shown in Fig. 4a,b, samples with input concentrations of 1 ng/μL yielded the highest fluorescent MBs ratio, while lower concentrations produced reduced ratios. In contrast, the lowest concentration (10^-4^ ng/μL), together with the negative control GrF3 (1 ng/μL) and the no-template control (NTC) groups, exhibited almost no fluorescent beads. Importantly, by applying the same fluorescence ratio threshold of 0.05, ASBVd positive samples were reliably distinguished across the 1-10^-3^ ng/μL range, but not at lower concentrations. This lowest detectable concentration (10^-3^ ng/μL) corresponds to 1.68 copies/μL, confirming the high sensitivity of our method.

**Fig. 4:**
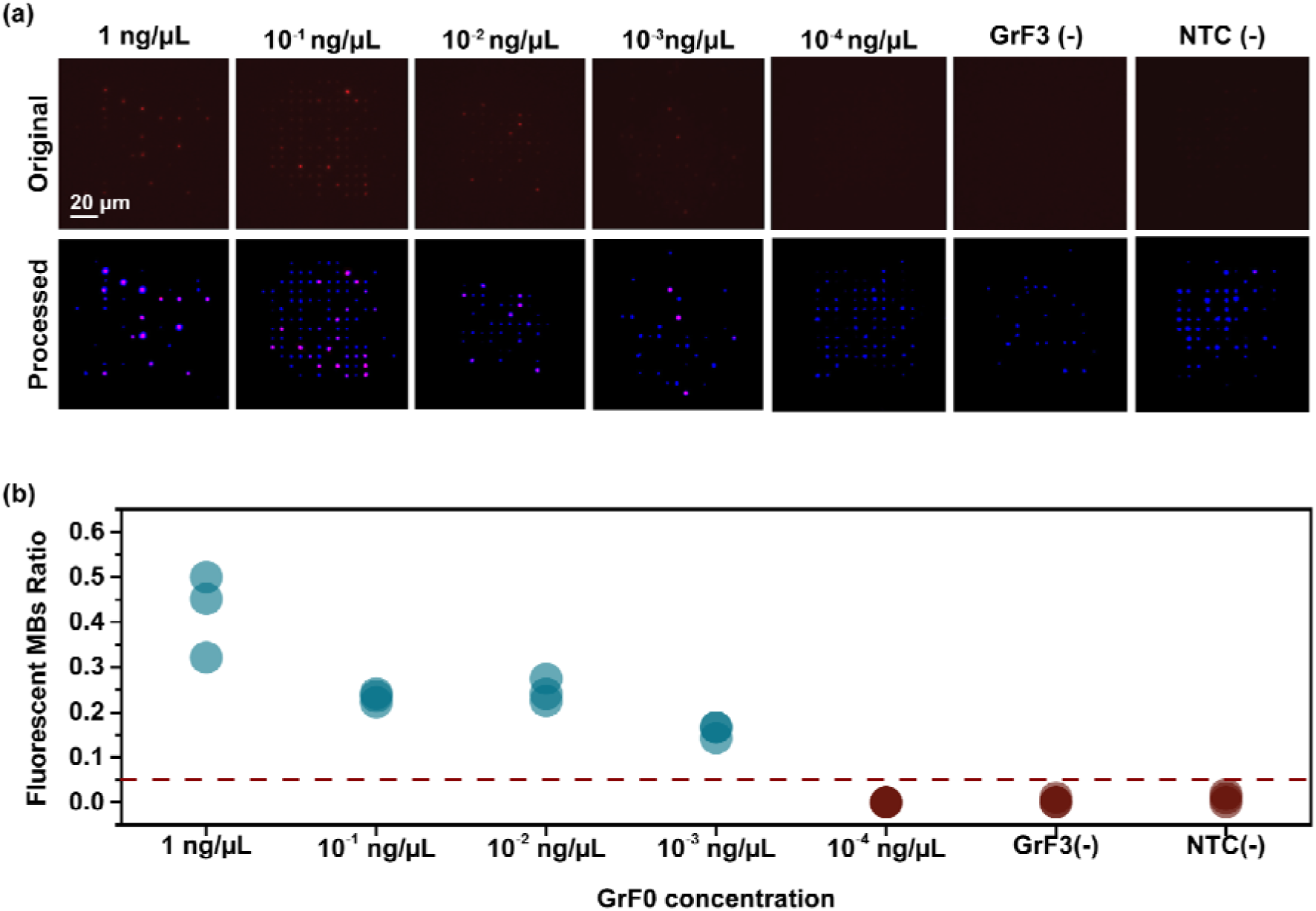
Nanopore array–based detection of GrF0 samples at different concentrations. **a** Images of nanopore array results for different concentrations of GrF0 samples. For each sample, the original image in the first row shows the raw fluorescent image captured by fluorescence microscopy. The processed image displays only the identified MBs in blue, and fluorescent MB-Cy5 in pink against a black background. **b** Detection results for different concentrations of the GrF0 sample, along with two negative samples (GrF3 with 1 ng/μL and NTC as a no-template control) for reference. Each condition is repeated on 3 different nanopore arrays.

Previously, Morey-León *et al.* reported a real-time reverse transcriptase polymerase chain reaction (RT-qPCR) assay for ASBVd with a detection limit of 8.8 copies/μL^26^. In comparison, our platform achieves a lower detection limit of 1.68 copies/μL, which is within one order of magnitude of ddPCR^18^, and more sensitive than RT-qPCR^26^. While our nanopore array does not provide linear quantification of viral copy numbers, this is inconsequential for practical plant screening, where binary determination of viral presence is sufficient. By establishing an appropriate fluorescent threshold, the nanopore array platform reliably distinguishes positive from negative samples, fully meeting the decision-making requirements of plant disease management protocols where timely intervention matters more than precise quantification.

For comparison, the CRISPR product of the GrF0 diluted samples were also tested using a fluorescence plate reader (see results in Fig. S2), where the result exhibited the same sensitivity of 10^-3^ ng/μL. It’s important to note that, for the plate reader, fluorescent signals plateaued at template concentrations ≥ 10^-2^ ng/μL, with no significant increase across different inputs. The observed signal plateau likely arises from several factors. During the RPA reaction, excess template can deplete critical cofactors, including recombinase proteins and single-stranded DNA-binding proteins, causing primers to compete inefficiently for these resources and to form non-productive primer-template complexes^27^. Meanwhile, high cDNA concentrations from the reverse transcription step can introduce inhibitory components such as excess dNTPs, residual RNA, and RT enzyme that interfere with downstream amplification^28^. In addition, the hook effect, where extremely high amplicon concentrations saturate Cas12a-crRNA complexes separately rather than forming productive ternary complexes necessary for trans-cleavage^29^, may further contribute to the signal plateau in the CRISPR detection step. Nevertheless, despite these saturation effects, such behavior benefits field applications by ensuring consistent positive detection across a wide concentration range.

Notably, the nanopore array readout did not perfectly mirror the bulk plate reader results, as the 1 ng/μL sample possesses a higher fluorescence ratio than other positive samples. This discrepancy can be attributed to two main factors: (i) variability in fluorophore conjugation efficiency and uniformity across assays, and (ii) differences in sampling scale—the nanopore array readout is based on the analysis of only tens to a few hundred beads per experiment, whereas the plate reader measures fluorescence from orders of magnitude more reporter groups in bulk. As a result, the nanopore array shows greater fluctuations by detecting signals from individual fluorescent beads. Nevertheless, when applied as a binary diagnostic tool to distinguish positive from negative samples, our nanopore array platform remains highly accurate and sufficiently sensitive.

Overall, our method consistently detected ASBVd at concentrations as low as 10^-3^ ng/μL (1.68 copies/μL), a sensitivity approaching that of ddPCR and exceeding that of RT-qPCR-based molecular diagnostics. These results highlight the potential of our nanopore array platform for ultra-sensitive, low-input viroid detection, which is further underscored by its successful detection of the viroid in all avocado tissues tested, such as leaf, flower, and fruit.

Beyond sensitivity and low input, this newly developed method also offers strong potential for portability in field use. Unlike conventional PCR and ddPCR platforms that require sophisticated thermal cycling and specialized laboratory infrastructure, our isothermal assay can operate with simple heating devices or even body heat supplemented with warming patches, making it particularly suitable for resource-limited field conditions. Among available isothermal amplification strategies, Loop-mediated isothermal amplification requires complex multi-primer design, while Helicase-Dependent Amplification suffers from slow kinetics. In contrast, RPA used in our method employs an elegantly simplified dual-primer design, facilitating assay development while maintaining robust performance across diverse sample matrices.

In addition to RPA, it is worth mentioning that rolling circle amplification (RCA) represents another promising isothermal method with potential for extremely low detection limits through its exponential signal amplification and generation of long concatenated DNAs. However, RCA currently suffers from slower reaction kinetics and requires circular templates or additional ligation steps. Despite these current limitations, future iterations of our platform could explore RCA integration to potentially push detection limits even further, particularly for applications where ultimate detection limits outweigh reaction time constraints.

Besides potential improvement with the assay protocol, our detection method could also benefit from advanced miniaturized sample-handling techniques capable of precisely processing with nL- or pL-scale volumes, such as electrowetting on dielectric (EWOD) digital microfluidics or surface acoustic wave (SAW) based droplet generation^30–33^. Integrating these approaches would fully leverage the low sample requirements of our nanopore array detection platform, thereby reducing both the effort of real sample collection and the overall reagent cost. Furthermore, while our current laboratory setup employs a compressed-air regulator for pressure-driven bead capture, future field-deployable versions could utilize simple manual pressure sources such as syringes or squeeze bulbs. On the imaging side, integration with compact CMOS-based fluorescence modules could further enable a miniaturized and portable detection platform. Together, these features highlight the promise of our approach as a highly sensitive and field-deployable diagnostic tool.

## Conclusion

In this study, we present a sensitive and low-input detection platform for Avocado Sunblotch Viroid (ASBVd) that integrates isothermal amplification, CRISPR-Cas12a–mediated signal conversion, and nanopore array–based single-particle analysis. The system demonstrated reliable performance across samples collected from multiple orchards and tissue types, yielding accurate positive and negative results. Compared with dLAMP assays reported in previous studies, our method exhibited higher sensitivity, detecting concentrations down to 10^-3^ ng/μL total cDNA (1.68 copies/μL). Furthermore, each readout consumes only 40 nL of the magnetic bead–fluorophore mixture from the entire CRISPR product, which is over 100-fold less than conventional fluorescence-based assays using plate readers. Unlike PCR-based approaches, our assay operates under isothermal conditions without the need for precise thermal cycling. Thereby, it reduces equipment requirements and enables rapid and portable detection, which is well-suited for field applications. With further integration and closed-cartridge development, this platform has the potential to be scaled for large-scale surveillance and adapted to detect a wide range of pathogens beyond ASBVd.

## Materials and Methods

### Sampling from avocado orchards in California

A total of 15 samples were selected from avocado orchards across Southern California for molecular analysis. The samples represented diverse tissue types including leaves, fruits, and flowers. Sample selection encompassed various infection states to evaluate pathogen detection capabilities, with tissues collected from trees displaying characteristic disease symptoms as well as neighboring asymptomatic trees. Pooled leaf, fruit, and flower samples were flash-frozen in liquid nitrogen and ground into a fine powder using a precooled mortar and pestle. The ground material was stored at −80°C until RNA extraction.

### RNA extraction and complementary DNA (cDNA) synthesis

Avocado tissue was homogenized in CTAB extraction buffer, extracted twice with chloroform/isoamyl alcohol (24:1, vol/vol), and precipitated with 10 M LiCl (final concentration, 3 M) overnight at −20°C. After ethanol washes, RNA was resuspended in nuclease-free water^18^.

The RNA was then purified using the Zymo RNA Clean & Concentrator-25 kit (Zymo Research, U.S.A.). RNA quality and concentration were evaluated using an Infinite Pro M200 NanoQuant (TECAN, Austria) and confirmed by agarose gel electrophoresis. cDNA was synthesized from total RNA using the HighCapacity cDNA Reverse Transcription Kit (Applied Biosystems, U.S.A.) performed at 37°C for 60 min, followed by 95°C for 5 min. The ASBVd copy numbers in the synthesized cDNAs were quantified using ddPCR^18^.

### RPA amplification and CRISPR-Cas12a detection

TwistAmp^@^ Basic kit was purchased from TwistDx^TM^. The RPA primers, crRNA, LbCas12a, and fluorophore–quencher probes were all obtained from Integrated DNA Technologies, and detailed information about the synthetic oligonucleotides are listed in Table S1. The RPA primer sets were designed using the PrimerQuest^TM^ online tool based on the ASBVd genome sequence targeted for amplification. Additionally, NEBuffer^TM^ r2.1 was purchased from New England Biolabs. The RPA reaction was conducted as per the manufacturer’s instructions: A mixture of 29.5 μL of rehydration buffer, 11.2 μL of nuclease-free water, and 2.4 μL each of forward and reverse primers (10 μM) was added to the enzyme pellet. Then, 2 μL of cDNA was added and mixed to achieve a total volume of 47.5 μL. Finally, 2.5 μL of MgOAc (280 mM) was added to initiate the reaction. The reaction mixture was incubated at 39 °C for 20 min. Following the incubation, 2 μL of RPA amplicons were added to a pre-assembled CRISPR-Cas12a reaction mixture comprising 50 nM of LbCas12a, 62.5 nM of crRNA, 10× buffer, and 2.5 μM of ssDNA BC-Q probe, resulting in a final reaction volume of 20 μL. The reaction solution was incubated at 37 °C for 30 min. After the incubation, the Cy5 fluorescence was measured by an Agilent BioTek Cytation 5 imaging/microplate reader (excitation/emission=645/665 nm).

### MB-Cy5 conjugation formation

Dynabeads^TM^ MyOne^TM^ Streptavidin C1 (Thermo Fisher Scientific) were prepared during the CRISPR incubation period. The beads were first resuspended by vortexing for 40 seconds, and 20 μL was transferred to individual tubes for each experimental group. 20 μL of 1× PBS buffer (Gibco^TM^, pH 7.4) was added to resuspend the beads, followed by magnetic separation using a NEBNext magnetic separation rack (New England BioLabs, Inc.) for 1 minute. After discarding the supernatant, the beads were removed from the magnet and resuspended in 20 μL of PBS buffer. This washing procedure was repeated three times, and after the final wash, the beads were resuspended in a PBS buffer. After characterization, the CRISPR products were transferred from plates to microcentrifuge tubes. An equal volume (20 μL) of pre-washed Dynabeads^TM^ MyOne^TM^ Streptavidin C1 was added to each tube. The mixture was incubated for 15 minutes at room temperature with gentle rotation. Subsequently, the biotinylated DNA-coated beads were separated using magnetic separation for 3 minutes. The coated beads were washed 3 times with a 1× PBS buffer to remove unbound material. Finally, the MB-Cy5 conjugates were resuspended in PBS buffer for TEM imaging and nanopore array–based single-particle fluorescence analysis applications.

### TEM Characterization

A JEOL 2100 transmission electron microscope (TEM) operating at 200 kV was used to capture images of the samples. An aqueous solution of magnetic beads, with or without DNA was deposited onto a carbon-coated copper TEM grid and allowed air dry. The dried grids were then rinsed in 18 Mega Ohm water for 30 s to remove excess salts and images were captured using a Gatan Orius camera.

### Nanopore array fabrication

A 200 × 200 µm^2^ suspended silicon nitride membrane with a thickness of 50 nm was fabricated via photo-lithography followed by KOH etching. A 15×15 array of nanopores (400 nm diameter) was drilled at the center of the membrane using a focused ion beam. The spacing between adjacent nanopores was 7 μm.

### Blockage-based detection set up

The nanopore array chip was mounted between two custom-fabricated chambers. The bottom chamber contained the sample solution with MBs or Cy5-MBs, while the top chamber with deionized water was exposed to ambient air. MBs or Cy5-MBs were diluted 1.25 × 10L-fold in deionized water to a final concentration of 580 aM. During the blockage experiment, a constant pressure of 0.5 atm was applied to the bottom chamber for 1 minute to drive bead capture. This external pressure is provided by compressed air through a pressure transducer (Type 3110 Pressure Transducer, Marsh Bellofram). Fluorescent images were then acquired. Details of the chip holder design are provided in a previous study^25^.

### Image capture and processing

All fluorescent images were acquired by an Olympus IX-81 inverted microscope with a 200 ms exposure time under the Cy5 channel. During the bead blocking, time-lapse fluorescence images were acquired every 5 s for 1 min to monitor the progression of bead capture and were used for the quantification of capture frequency. For the fluorescence-ratio analysis reported in the main text, only the image acquired after 1 min of pressure-assisted bead capture was used. Image processing and nanoparticle counting were performed using ImageJ software. Before thresholding, a Gaussian blur filter (sigma = 2.0) was applied to all fluorescence images. An intensity threshold of 160 was applied to non-biological sample demonstration (Fig. 2), while a threshold of 180 was used for all real-world avocado samples (Fig. 3-4).

## Acknowledgements

This work is supported by USDA-NIFA under award number 2024-67014-42672. K.D. acknowledges support from the National Institutes of Health under award number R35GM142763. J.X., Y. Z., and C.D. would like to thank the Photonics Center at Boston University and the Center for Nanoscale Systems at Harvard University for providing access to their fabrication and characterization facilities.

## Author Contributions

J.X., X.J., and M.K.D. designed and performed the experiments. J.X. and X.J. drafted the manuscript. Y.Z. contributed to device fabrication. B.S. participated in the experimental design and manuscript revision. R.P. assisted in the experiments. C.D., F.K., and K.D. secured funding and supervised the project and revised the manuscript. All authors discussed the results and approved the final version of the manuscript.

## Competing Financial Interests statement

The authors declare no competing financial interests.

## Notes

### Competing Interest Statement

The authors have declared no competing interest.

