## Supplementary Information for "Ultrasensitive, Low-Input Detection of Avocado Sunblotch Viroid via RPA-CRISPR and Nanopore-Array Single-Bead Fluorescence Readout"

### Supporting Information

**Table S1**: Description of collected avocado samples used in this study. The number after the location specifies the number of orchards.

| Sample Name[a] | Location[b] | Tissue | Tree code[c] | Copy number/µL (ddPCR) |
|---|---|---|---|---|
| GrF0 | San Diego 1 | Fruit | 0 | 1919 |
| GrL0 | San Diego 1 | Leaf | 0 | 304 |
| GrFW0 | San Diego 1 | Flower | 0 | 704.3 |
| GrL1 | San Diego 2 | Leaf | 1 | 427.2 |
| GrF1 | San Diego 2 | Fruit | 1 | 398 |
| GrL3 | San Diego 2 | Leaf | 3 | 0 |
| GrF3 | San Diego 2 | Fruit | 3 | 0 |
| OGL1 | San Diego 3 | Leaf | 1 | 43.76 |
| JsL2 | San Diego 4 | Leaf | 2 | 2.4 |
| AlL0 | Ventura | Leaf | 0 | 0 |
| AlF0 | Ventura | Fruit | 0 | 0 |
| AlL10 | Ventura | Leaf | 10 | 0 |
| AlF10 | Ventura | Fruit | 10 | 0 |
| AcL7 | Riverside | Leaf | 7 | 0 |
| AcF7 | Riverside | Fruit | 7 | 0 |

[a] The first letters in the label correspond to the names of the orchards.

[b] The number after the location specifies the orchards code.

[c] The number refers to the specific tree. As not all samples from the previous publication were checked here, some numbers are missing.

**Table S2**: Sequences of synthetic Oligos used in this study.

| Name | Sequence (5’-3’) |
|---|---|
| crRNA1 | ACGAAACCAGGTCTGTTCCGA |
| Primer Set 1 Forward | AGAACAAGAAGTGAGGATATGATTAAACTTTGTTTG |
| Primer Set 1 Reverse | CGAAGTGATCAAGAGATTGAAGACGAGTGAAC |
| crRNA2 | CCTGAAGAGACGAAGTGATCA |
| Primer Set 2 Forward | GACTCTGAGTTTCGACTTGTGAGAGAAGGA |
| Primer Set 2 Reverse | CAATGAAGATAGAGGAGTAAACCTTGCGAGAC |
| ssDNA BC-Q probe | /5BiosG/iCy5/TTATT/3IAbRQSp/ |
| ssDNA positive control | /5BiosG/iCy5/TTATT |

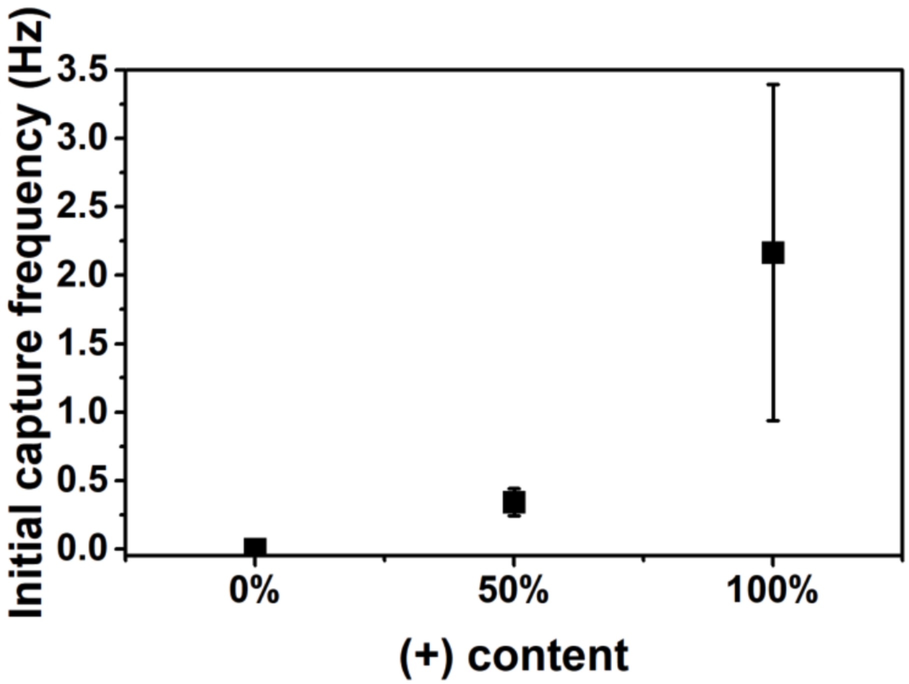


**Fig. S1**. Initial capture frequency analysis for different MB-Cy5 contents. The rate of MB–Cy5 blockage events decreased with lower proportions of MB–Cy5, consistent with the correlation between capture frequency and fluorescent nanoparticle concentration.

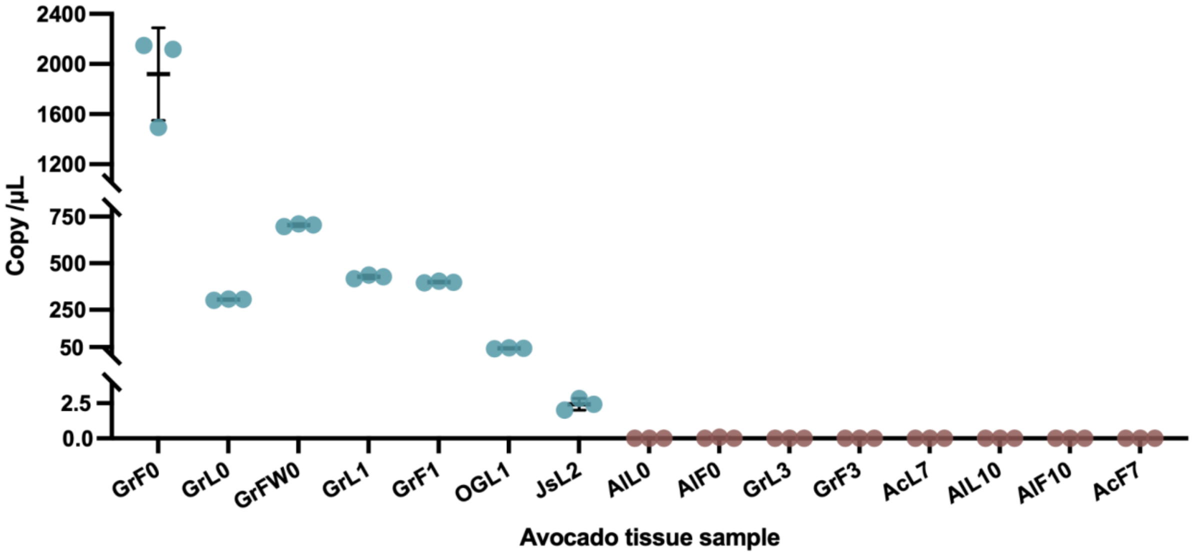


**Fig. S2**. ddPCR result of ASBVd number count for different avocado tissue samples. Negative samples for the number count can not be detected are marked in red.

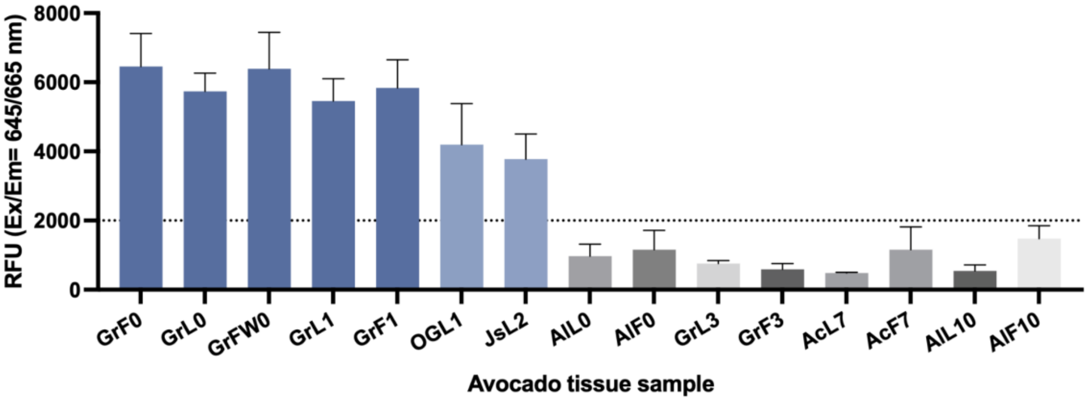


**Fig. S3**. Specificity results from the image reader (excitation/emission=645/665 nm).

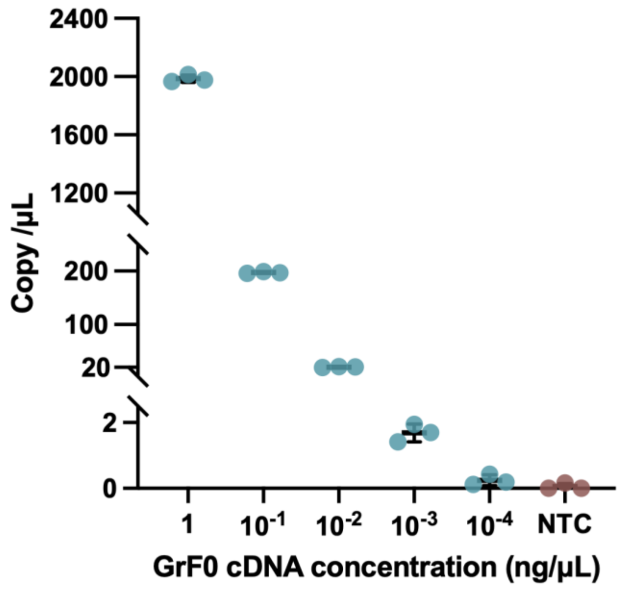


**Fig. S4**. ddPCR result of ASBVd number count for different concentrations for GrF0 sample along with negative controls. Negative samples for the number count can not be detected are marked in red.

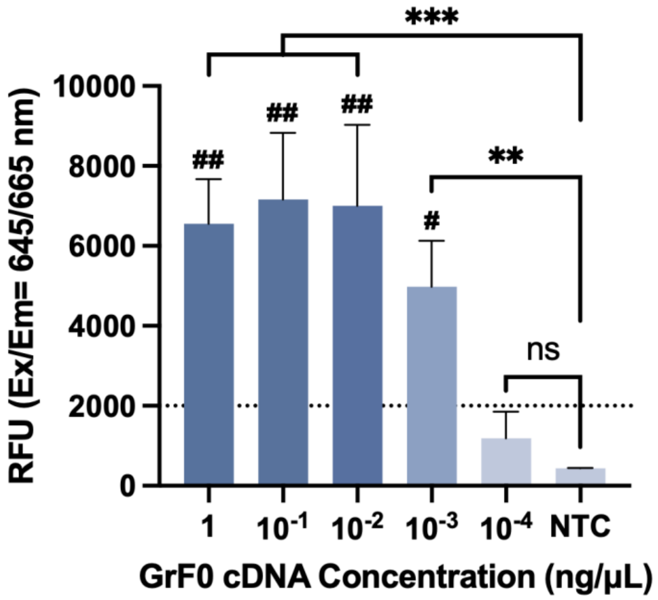


**Fig. S5**. Sensitivity results from the image reader (excitation/emission=645/665 nm).

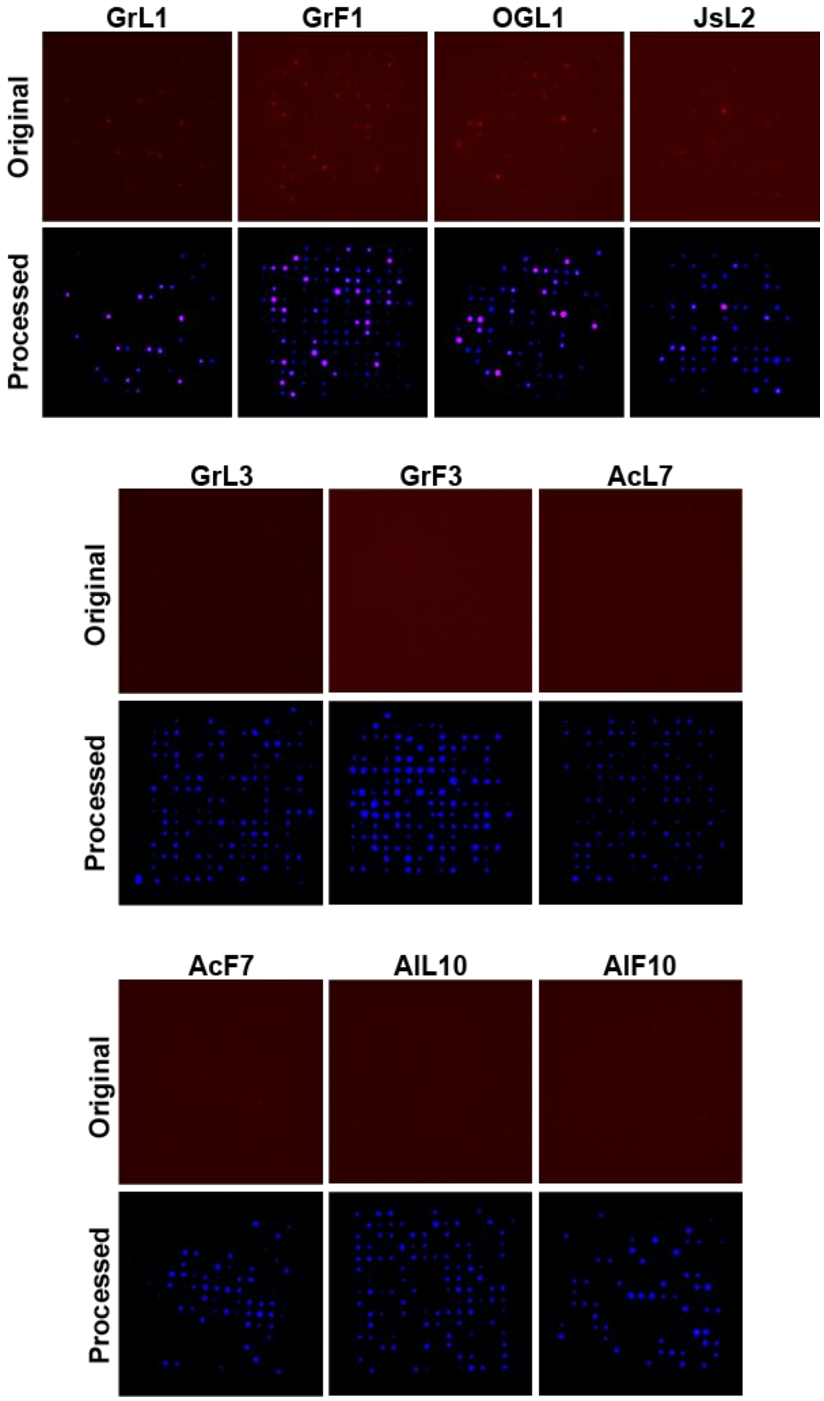


**Fig. S6**. Detection results of individual collected avocado samples. For each sample, the original fluorescent image shows the raw signal captured by fluorescence microscopy. The processed (highlighted) image displays only the identified magnetic beads (MBs) in blue, and fluorescent MBs in pink against a black background. The 4 samples in the first row are identified as positive samples from PCR results; the other six samples are negative.

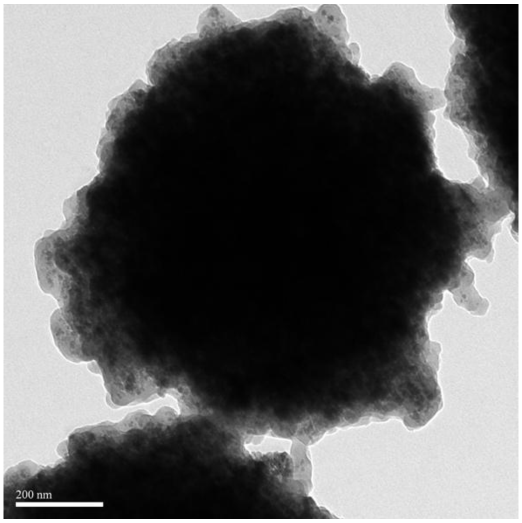


**Fig. S7**. Transmission electron microscopy characterization of magnetic beads. TEM image of magnetic beads used in the detection system, showing characteristic spherical morphology with accessible surface area suitable for biomolecular conjugation.
